# Investigating the sensitivity of the diffusion MRI signal to magnetization transfer and permeability via Monte-Carlo simulations

**DOI:** 10.1101/2025.07.16.664944

**Authors:** Zhiyu Zheng, Karla L. Miller, Benjamin C. Tendler, Michiel Cottaar

**Affiliations:** Centre for Integrative Neuroimaging, FMRIB, Nuffield Department of Clinical Neurosciences, University of Oxford, Oxford, United Kingdom

## Abstract

**Purpose:** Magnetization transfer (MT) and water exchange via permeability operate on a similar spatiotemporal scale to water diffusion. In this study, we use a simulation-based approach to characterise how MT and permeability impact (1) diffusion-weighted MRI (dMRI) measurements from cylindrical substrates and (2) parameter estimation using a two-compartment model of white matter.

**Methods:** We used Monte-Carlo simulations to model the dMRI signal inside and outside axons by simulating signals from parallel cylinders with different diameters and volume densities. We subsequently introduced membrane permeability and MT at the cylinder walls to investigate their impact on the dMRI signal. We fitted a two-compartment model to the simulated signal to produce estimates of the cylinder diameter and density. We evaluated the impact of MT and permeability by comparing the fitted diameter and density to the simulated ground truth.

**Results:** Permeability leads to underestimation (up to 100%) of cylinder diameter and density. Specifically, by enabling isochromats to escape from restrictions and diffuse more freely, permeability makes the overall displacement profile closer to the extra-axonal displacement profile. MT had limited effects on diameter estimation but caused substantial bias (20-50%) in volume density estimates depending on the ratio of the intra-axonal and extra-axonal volume fraction. This is due to the intra-axonal and extra-axonal space having different surface-to-volume ratios and therefore different surface relaxation rates.

**Conclusion:** Permeability and MT can considerably influence the dMRI signal. They increase the relative contribution from larger cylinders to the dMRI signal and bias microstructural parameter estimates derived from dMRI data.

## 1. Introduction

Diffusion-weighted MRI (dMRI) is a leading technique for the non-invasive characterization of tissue microstructure. Whilst MRI methods are typically associated with imaging voxels on the order of a cubic millimeter, dMRI yields image contrast driven by the diffusion of water in tissue on the order of micrometers. Measured signals are sensitive to changes in cellular morphology and density, providing a basis to characterize brain microstructure, composition, and changes with disease.

To estimate the cellular properties of tissue from diffusion MRI, microstructure models are fit to experimental data acquired in multiple diffusion regimes (e.g., with different gradient orientations/durations/amplitudes/waveforms or diffusion times). Typically, these models condense a complicated tissue environment into a few distinct compartments corresponding to key tissue features. For example, a widely used signal model for characterizing white matter is AxCaliber^1–3^. It models the dMRI signal as two compartments corresponding to intra-axonal and extra-axonal water. The intra-axonal compartment is modeled as parallel cylinders with finite diameter, and the extra-axonal compartment is modeled as a Gaussian diffusion tensor.

AxCaliber provides an estimate of two tissue properties that have attracted considerable research interest: axon diameter and axon density (volume fraction). Axon diameter has been reported to correlate with neuron’s conduction velocity^4,5^ and myelination level^6^. Recently, axon diameter mapping has also been used to study the development of the human nervous system non-invasively^7,8^. Axon density is also closely linked to neurodegenerative diseases as it can (indirectly) reflect pathological cell loss or neuron regeneration^9–11^.

A common feature of AxCaliber and other multi-compartment dMRI signal models is the assumption that signal attenuation (diffusion-weighted signals relative to b=0) is driven solely by diffusion effects. However, it is well established that several other effects occur at similar spatiotemporal scales to diffusion, including water exchange across semipermeable membranes^12^ and magnetization transfer (MT)^13^. Specifically, membrane permeability arises from water passing through cross-membrane transport proteins (e.g., aquaporins) and/or osmosis^14^. It allows magnetization carried by water molecules and other particles to enter and escape from compartments, leading to a further change in displacement profiles and signal amplitude beyond diffusion-related changes^12,15–18^. MT describes the transfer of magnetization between different molecules. MT is of most interest when magnetization transfers between a free water molecule and a macromolecule, the latter being characterized by very short transverse relaxation times (<1ms^19,20^). Magnetization that has transferred from water molecules into a macromolecule will typically vanish before it can be measured with MRI, reducing the signal amplitude. Since most current models attribute all signal attenuation to diffusion, MT and permeability-induced signal attenuation is a potential source of bias to fitted model parameters. For example, these effects could contribute to the bias observed between axon diameter estimates from dMRI and histology-derived estimates^21–29^.

To date, whilst there are established experimental methods to investigate permeability in tissue using MRI^30–34^, parameter estimation is typically performed using analytical models consisting of multiple Gaussian compartments. These models do not incorporate time-dependent diffusion properties arising from geometries of restrictions (preventing the estimation of axon diameter), and cannot be easily translated to more realistic tissue models^17,35,36^. Proposed analytical solutions also cannot be easily translated to more realistic tissue geometries. As for MT, no analytical solutions exist when considering dMRI signals.

One approach to investigate the impact of permeability and MT on the dMRI signal is via Monte-Carlo simulations, which can incorporate tissue models of arbitrary complexity. In this project, we use Monte-Carlo simulations to investigate how the estimation of microstructural tissue features in models of white matter from dMRI are impacted by the contribution of permeability and MT. We performed our investigation using a novel Monte-Carlo simulator developed by our research group, MCMRSimulator^37^, that can incorporate the effects of diffusion, permeability, MT and off-resonance field perturbations.

Specifically, we simulated the dMRI signal generated by isochromats (i.e., an ensemble of spins experiencing the same off-resonance field) in substrates consisting of randomly distributed parallel cylinders. Parallel cylinders is a commonly used approximation for axons in white matter and our simulations included both fixed and distributed diameters and incorporated different levels of permeability and MT. We subsequently fit a two-compartment model to the signal to estimate the axon diameter and density and evaluated the parametric estimation bias introduced by permeability and MT.

We found that the impact of permeability and MT depends on the specific axon diameters and their distribution. When axons were modelled as uniform diameter cylinders, permeability led to underestimation of both axon diameter and density. MT led to an over or underestimation of axon density (dependent on the underlying intra-axonal volume fraction), but little bias on diameter estimates. Importantly, we demonstrate that a small amount of permeability (exchange time ~600ms) is enough to make a two-compartment model insensitive to axon diameters below 2μm, corresponding to realistic axon sizes in the central nervous system^38^. When axons were modelled as cylinders with a distribution of diameters, permeability still led to an underestimation in both axon diameter and density, but MT introduced an overestimation in both axon diameter and density. We discuss how these observations may be generalizable beyond white matter to a wide range of dMRI signal models and substrates, with software provided for readers to replicate our simulations and perform their own investigations.

## 2. Methods

### 2.1 Monte-Carlo Simulation

All simulations were performed using MCMRSimulator v0.9^37^, a Julia language package developed for modelling the impact of diffusion, relaxation, MT, permeability and off-resonance on different MRI sequences and microstructural substrates. Below we describe the implementation of permeability and MT with MCMRSimulator, and the subsequent simulation study.

#### 2.1.1 General simulation parameters

##### 2.1.1.1 Substrate

All experiments simulated isochromats diffusing inside randomly distributed parallel cylinders and the free space outside the cylinders. Figure 1 shows two example substrates with (a) uniform and (b) distributed diameters. The cylinder walls were modelled as having zero thickness. Two sets of simulations were performed: 1. All cylinders have a uniform diameter, with the diameter varying from 0.6 to 11μm across different substrates (fixed diameter case); 2. Cylinders have a distribution of diameters corresponding to a Gamma distribution with mean of 4μm and variance of 1μm^2^ (distributed diameter case). These diameter range and distribution was chosen based on previous microscopy and MRI studies^22,29,39,40^. For these simulations, the volume fraction inside the cylinders were set to 0.65 to approximate realistic axon densities in white matter^41–44^. The cylinder packing was performed by randomly distributing the number of cylinders needed to achieve the given density and then using distance-dependent repulsion between cylinders to eliminate overlaps among them and create a near-homogeneous spatial distribution.

**Figure 1.**
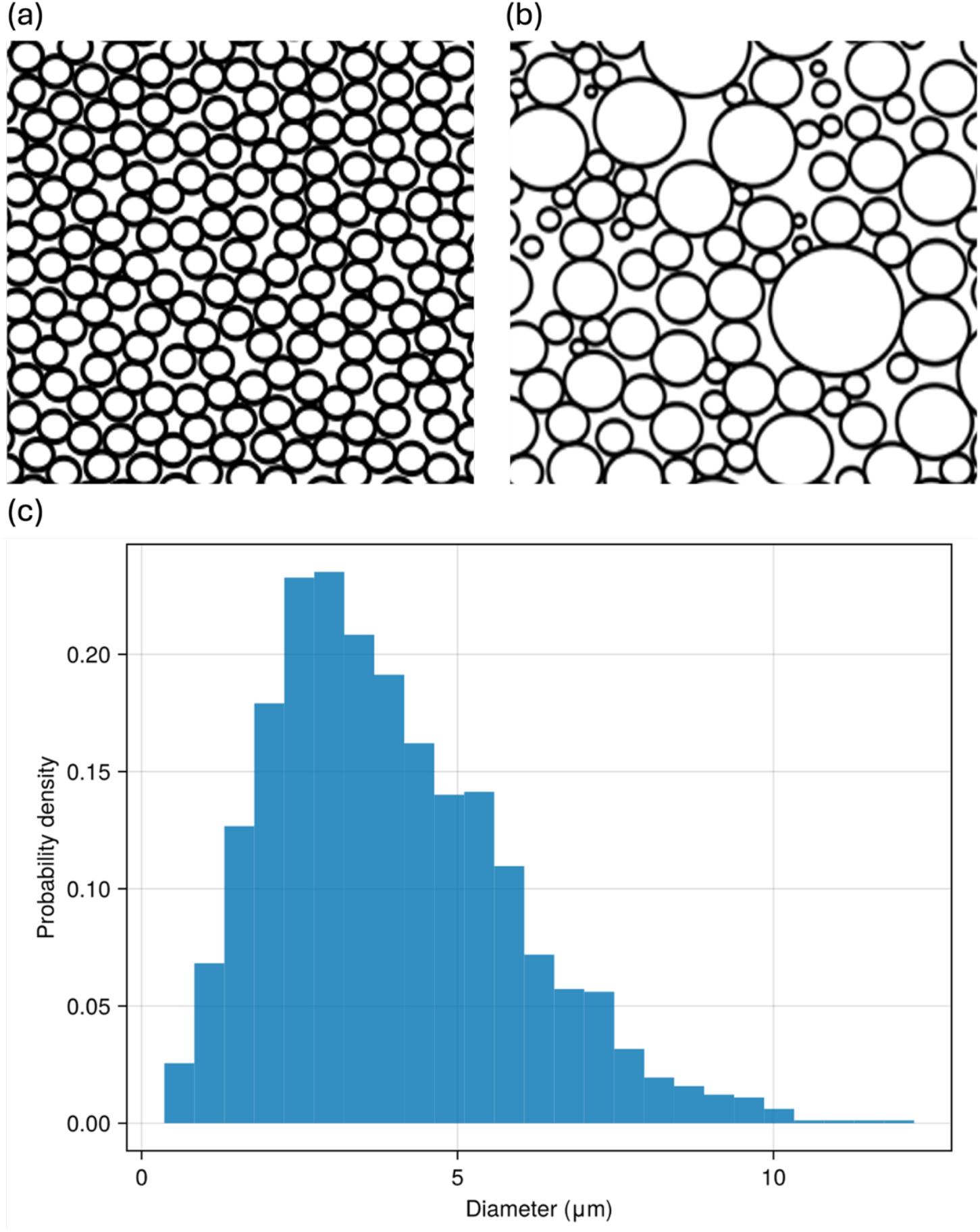
Investigated Monte-Carlo substrates. Figures 1a and b display the simulated substrates consisting of (1) randomly distributed cylinders with an identical diameter (Figure 1a) and (2) randomly distributed cylinders with Gamma distributed diameters (Figure 1b). The Gamma distribution corresponds to a mean of 4μm and standard deviation of 1μm^2^ (Figure 1c). Default simulations were performed with a fixed intra-cylinder volume fraction of 0.65.

##### 2.1.1.2 Sequence parameters

We used the original AxCaliber pulsed gradient spin echo sequence parameters for our simulations^1^. Specifically, we adopt a gradient duration (δ) of 2.5ms; gradient strengths incremented from 0 to 1200mT/m with steps of 80mT/m; diffusion times (Δ) of {10, 15, 20, 30, 40, 50, 60, 80}ms; TE of 166ms. The diffusion-weighting gradients were applied perpendicular to the principal axis of the cylinder.

##### 2.1.1.3 Monte-Carlo simulation setup

Even within a 1mm^3^ voxel there could be more than 10^20^ spins which are computationally unrealistic to simulate. Therefore, here we simulated a random subset of isochromats – magnetization representing an ensemble of spins that have similar trajectories and, hence, experience the same off-resonance field. 250,000 isochromats were simulated in each experiment to achieve a good balance of accuracy and simulation speed. The isochromats were initially randomly distributed over an isotropic voxel with its side length equal to 100 times the mean cylinder diameter. For all isochromats, the intrinsic diffusivity was set to 2.3μm^2^/ms to approximate the intra-axonal axial diffusivity previously reported in human tissue^45^. The intrinsic relaxation times (T_1_, T_2_) were set to infinity as they equally impact the diffusion and non-diffusion weighted signal. MT introduces effective T_2_ difference between compartments, which we will discuss in 2.1.2.2.

#### 2.1.2 Implementation of MT and permeability

##### 2.1.2.1 Permeability

To model permeability, we introduced a geometry-independent parameter to characterize the probability of an isochromat passing through an obstruction for a simulation with a 1ms timestep. As shown in Figure 2a, the isochromat keeps its original direction of travel if it passes through the obstruction (dashed line) and gets elastically reflected otherwise (solid line). The permeability parameter sets the local probability of isochromats passing the membrane and indirectly controls the exchange time. It was varied across a wide range (0.001-0.02) to explore different membrane types and different membrane states.

**Figure 2.**
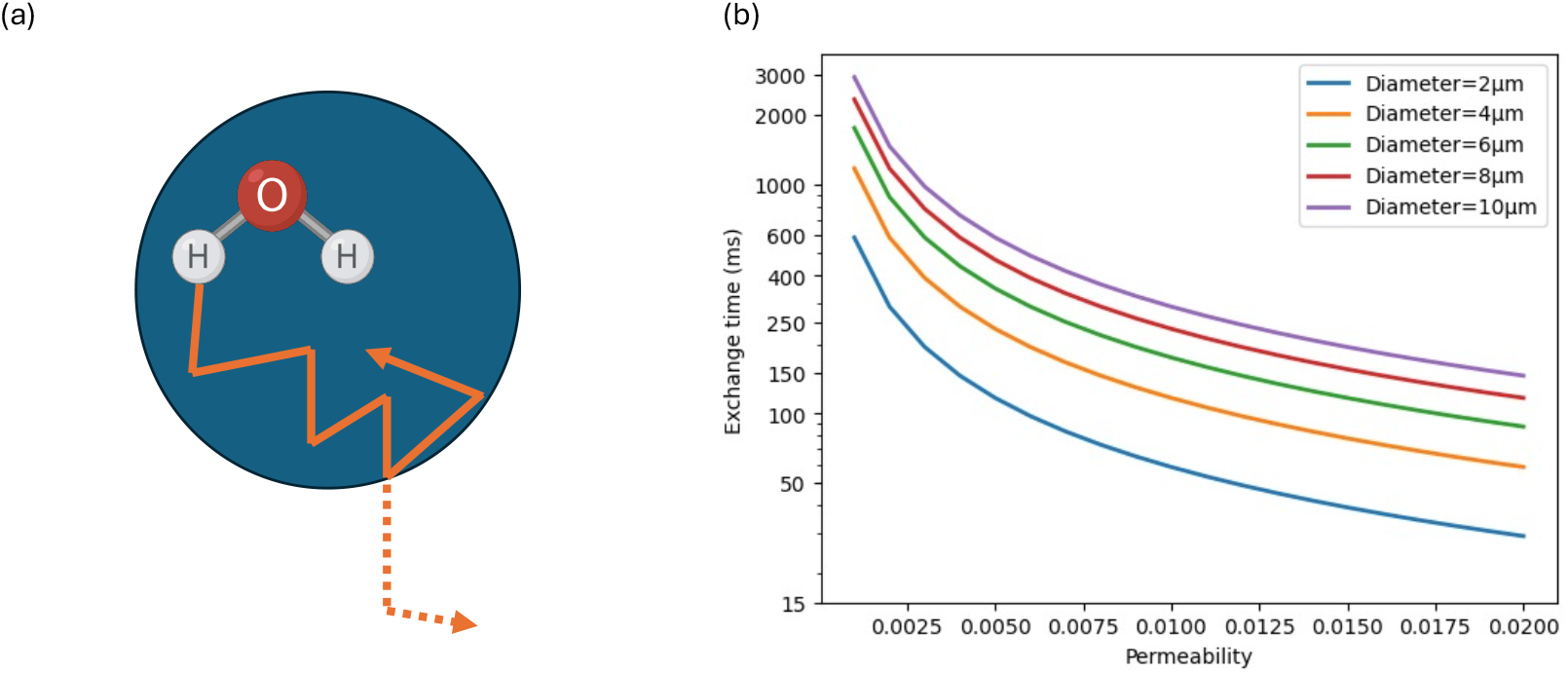
Permeability implementation: Figure 2a illustrates if a water molecule collides with an obstruction (cylinder wall), it has a probability of continuing in the original direction and crossing the membrane (dashed line), rather than being reflected (solid line). The probability of crossing is defined as the *permeability* parameter in the simulation. Figure 2b shows the mapping between *permeability* and exchange time at different cylinder diameters. Note that the exchange time depends on the diameter because opportunities for isochromats to exchange are driven in part by surface-to-volume ratio.

A common parameter to characterize membrane permeability is exchange time^17,46–49^. It is an exponential decay coefficient characterizing the average time it takes for a particle in one compartment to exchange to another compartment. The exchange process can be described by the following exponential relationship:

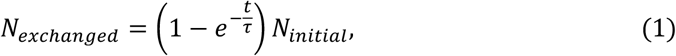

where τ is the exchange time constant, *N*_*exchanged*_ is the number of particles that have exchanged to another compartment, and *N*_*initial*_ is the number of particles initially inside the compartment.

The exchange time can be analytically derived from the rate of isochromats passing the surface, with previous work^37^ defining the exchange rate *R*:

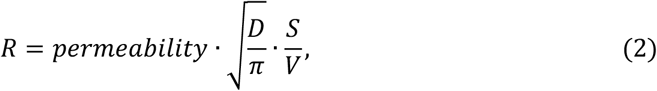

where *permeability* is the probability of an isochromat passing through an obstruction, D is the intrinsic diffusivity, S is the surface area between the two exchanging compartments and V is the total volume. S/V is the surface-to-volume ratio and for cylinders:

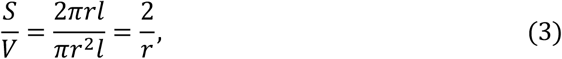

where r is the radius, l is the cylinder length. As the exchange time constant τ is the reciprocal of exchange rate R, we can combine Eq.2 and Eq.3 to have:

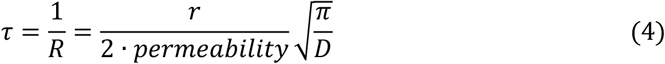

For a cylinder diameter of 2μm, our chosen range of permeabilities corresponds to exchange times of 29-584ms. As shown in Figure 2b, cylinder diameters are closely related to the exchange time. We describe how this relationship impacts parameter estimation in section 4.1.

##### 2.1.2.2 Magnetization Transfer (MT)

###### Modelling approach

In this work, we model MT as an exchange process at the membrane. Specifically, MT occurs when an isochromat collides with an obstruction (e.g. a cylinder wall). To model the interaction during the collision, a bound-pool interaction model was implemented. As shown in Figure 3, isochromats are divided into two groups of protons that exchange with each other – the bound pool and the free pool. The bound pool represents ensembles of protons on macromolecules localized at user-defined obstructions and has a very short transverse relaxation time, T_2_. The free pool represents ensembles of protons in water molecules and occupies the remaining space. When an isochromat encounters the obstruction, it has a certain probability of getting bound to the obstruction (i.e., transferred into the bound pool). Once bound, it will experience very short T_2_ and quickly loses all its transverse magnetization. After some time, it is released from the bound pool and subsequently takes a new random step which is consistent with the defined diffusion coefficient.

**Figure 3.**
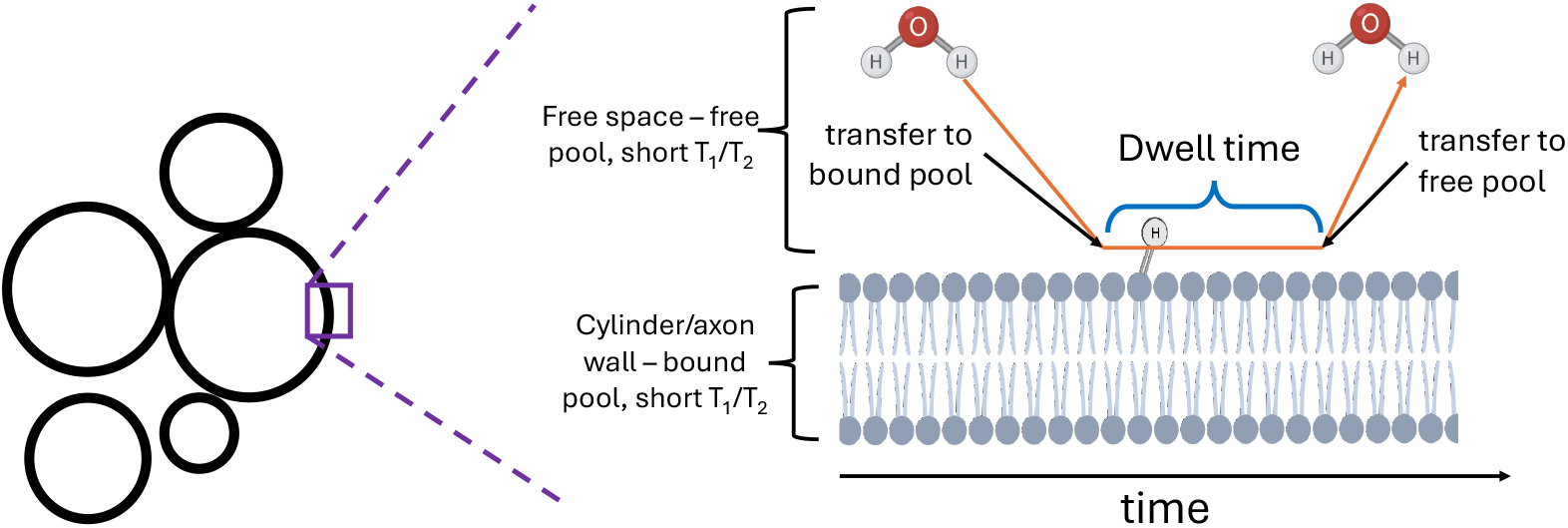
Illustration of the bound pool interaction mechanism at the cylinder wall: Isochromats are split into a free pool and bound pool. The bound pool is localised at the obstructions (membrane in this case). When an isochromat in the free pool encounters the membrane, there is a fixed probability that it will transfer into the bound pool and get spatially attached to the membrane. This is controlled by both surface density and dwell time. The dwell time is the characteristic timescale for a bounded isochromat to be transferred back into the free pool.

We control the properties of MT via two characteristics: (1) dwell time characterizes the average time an isochromat remains in the bound pool; (2) surface density is the ratio between surface isochromat density on the obstruction and the volume isochromat density in the free pool. Dwell time and surface density together control the probability of an isochromat getting bounded when it encounters an obstruction. The simulator uniformly distributes the isochromats in the two pools over their corresponding surface/volume during initialization. A more detailed description of the implementation can be found in the simulator paper^37^.

###### Parameter selection

Whilst the intrinsic relaxation constants of the free pool were set to infinity, MT introduces relaxation through the bound pool. Isochromats that transfer from the free pool to the bound pool quickly lose their transverse magnetization, reducing the total transverse magnetization of the whole system. As a result, MT effects can be observed and quantified in the b=0 signal as an effective T_2_ relaxation, where a shorter effective T_2_ means a stronger contribution of MT. Eq.5 below defines the relationship between dwell time, surface density and effective T_2_ for cylinders. See Appendix for a detailed derivation.

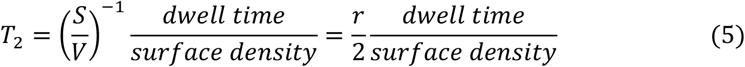

Since we only consider the signal within a single TR, there’s no saturation effect and MT’s impact on T_1_ can be ignored.

Appropriate bound pool parameters were set to achieve effective T_2_s at the same order of magnitude as the T_2_ of white matter (~80ms at 3T). The dwell time was set to 30ms as reported by Helms et al.^50^ and the surface density was varied to achieve different T_2_ values ranging from 30 to 150ms. The T_2_ of the bound pool (macromolecules) was set to 1μs, while the T_1_ remained infinite. For the distributed cylinder diameter case, the effective T_2_ is dependent on the cylinder radius due to the surface-to-volume ratio. Therefore, we varied the surface density to achieve different effective T_2_ for a cylinder diameter of 4μm (the mean diameter of the distribution) and used the same surface density for all the cylinders in the set.

### 2.2 Two-compartment model fitting

To estimate the cylinder diameter and density from a simulated signal, we fit a two-compartment model to our data. Specifically, we modelled the diffusion-weighted signal attenuation as the sum of two compartments corresponding to intra-axonal and extra-axonal water, defining:

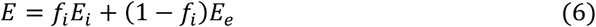

where *f*_*i*_ is the intra-axonal signal fraction that quantifies the cylinder density, *E*_*i*_ is the intra-axonal signal attenuation and *E*_*e*_ is the extra-axonal signal attenuation. We modelled the extra-axonal signal as a free diffusion compartment that exhibits Gaussian diffusion and uniform radial diffusivity perpendicular to the cylinder axis. Thus, the extra-axonal signal is represented by:

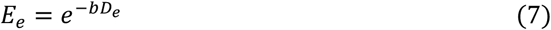

where *b* is the b-value and *D*_*e*_ is the diffusivity of the extra-axonal space. Note that due to hinderance of the cylinders *D*_*e*_ is not necessarily the intrinsic diffusivity and therefore will need to be estimated from the data.

For the intra-axonal signal, we used the MCMRSimulator to generate a dictionary of intra-axonal signals from parallel cylinders with different diameters at a spacing of 0.2μm. We subsequently constructed a projection from cylinder diameter to the intra-axonal signal by interpolating between the discrete value pairs in the simulated dictionary. This approach allowed us to model intra-axonal signals more accurately than existing analytical models based on Gaussian phase approximation^51,52^ (see supporting information 1.1). We then used these representations of the intra-axonal and extra-axonal signal attenuations to describe the total diffusion attenuation via:

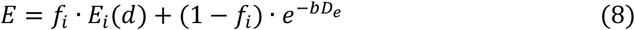

Fitting Eq.8 to a simulated dMRI signal allows us to estimate the cylinder density (i.e., intra-axonal signal fraction, *f*_*i*_), mean cylinder diameter (*d*), and extra-axonal diffusivity (*D*_*e*_).

Data analysis was performed with Python (version 3.11). Interpolation of the simulated dictionary was performed using the CubicSpline function in SciPy^53–55^. We used the minimize function from SciPy (Nelder-Mead algorithm) to fit our model to the simulated data and estimate three microstructural parameters: cylinder diameter, density and *D*_*e*_. The optimization was constrained with the following conditions to prevent unrealistic estimates: *d* ∈ [0,20]*μm, f*, ∈ [0,1], *D*_*e*_ ∈ [0,4]*μm*^2^/*ms*. The intra-axonal diffusivity was set to 2.3μm^2^/ms, same as the intrinsic diffusivity.

## 3. Results

All the scripts and raw data that are used to obtain the following results are available online at https://github.com/zhiyuzheng1769/MT-and-permeability-effect-on-two-compartment-dMRI-WM-model. As defined in 2.1.1.1 we present results from simulations using a substrate with uniform cylinder diameters (fixed diameter case) and distributed cylinder diameters (distributed diameter case).

### 3.1 No MT and Permeability

Figure 4a,b displays the estimated diameter and volume fraction as a function of the true underlying cylinder diameter for the fixed diameter case. We observed a small bias in the estimates for cylinder diameter and volume fraction as a function of cylinder diameter, with diameter estimate errors within ±1μm and <0.05 for the volume fraction estimation.

**Figure 4.**
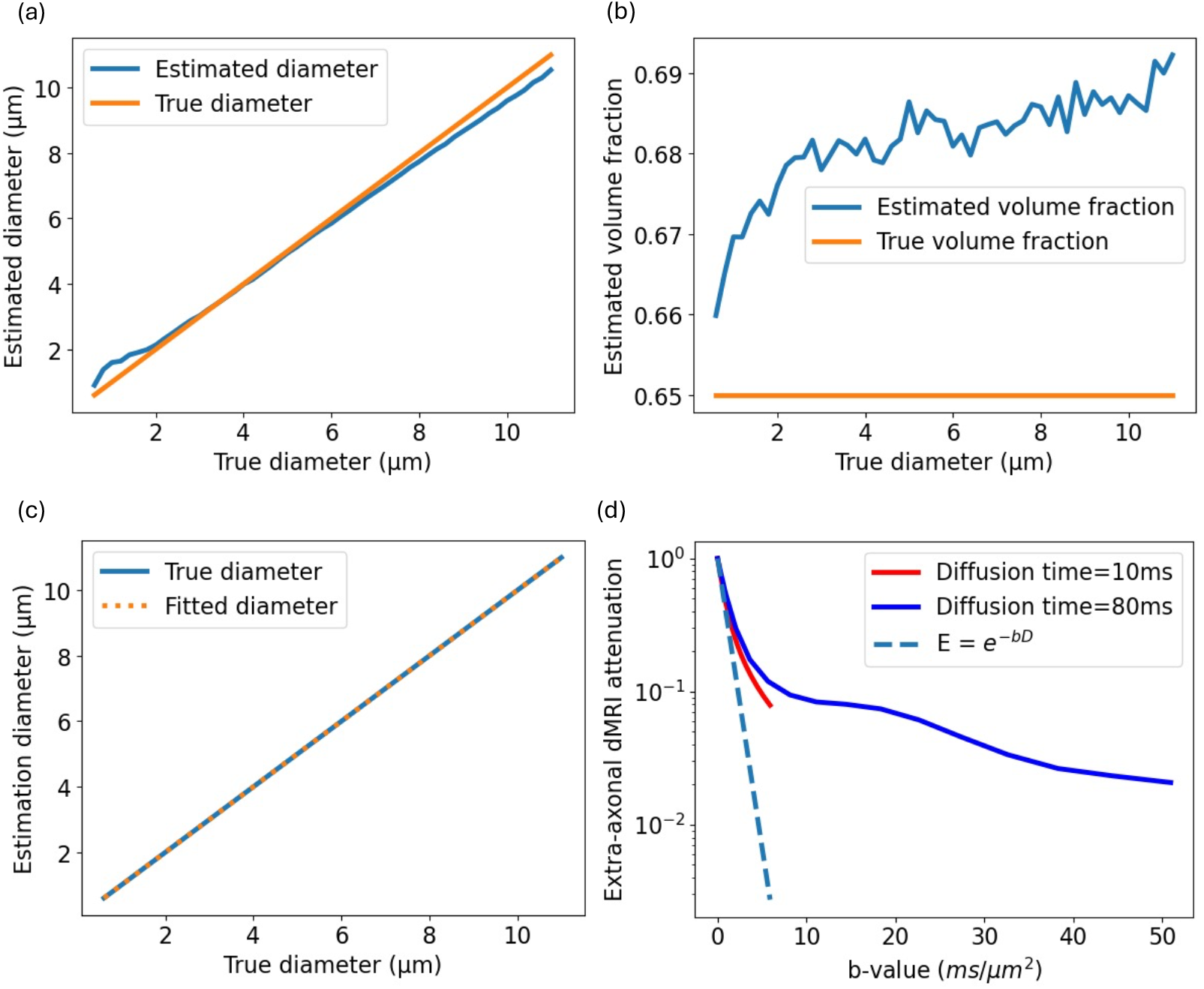
Parameter estimation and simulated signal with no permeability or MT (fixed diameter case). Figure 4a and 4b display the cylinder diameter and volume fraction estimates without permeability or MT. Figure 4c shows the cylinder diameter estimates based on a model including only the intra-axonal signal compartment, and Figure 4d shows simulated extra-axonal signal attenuation. The extra-axonal attenuation is not linear with respect to b-value on the semilog plot, inconsistent with a Gaussian diffusion model (*E* = *e*^−*bD*^).

Separating the intra-axonal and extra-axonal signals (Figure 4c,d), we observed that whilst the cylinder diameter can be accurately fitted by our simulation-based intra-axonal signal model, the extra-axonal signal compartment does not exhibit Gaussian diffusion behavior across all investigated diffusion regimes. Further investigation found that including higher order non-Gaussian diffusion terms (kurtosis or time-dependent diffusion) in the model of the extra-axonal signal did not improve the estimation accuracy (Supporting information 1.2). To account for the small bias in Figure 4a, all findings for the permeability and MT investigations are described relative to these estimates.

### 3.2 Permeability

Figure 5 displays the estimated diameter and volume fraction for semipermeable cylinders (fixed diameter case). The introduction of membrane permeability leads to diameter and volume fraction underestimation. For both cylinder diameter and volume fraction estimation, we observe that the bias relative to the no permeability case becomes larger as the true cylinder diameter decreases and as the permeability of the cylinder wall increases. Notably, there exists a cutoff diameter as a function of permeability, below which the signal model estimates zero diameter. Furthermore, we see that a low level of permeability -- 0.001, corresponding to an exchange time of 584ms for a 2μm-diameter cylinder, is already enough to cause visible underestimation in diameter and volume fraction for small cylinders.

**Figure 5.**
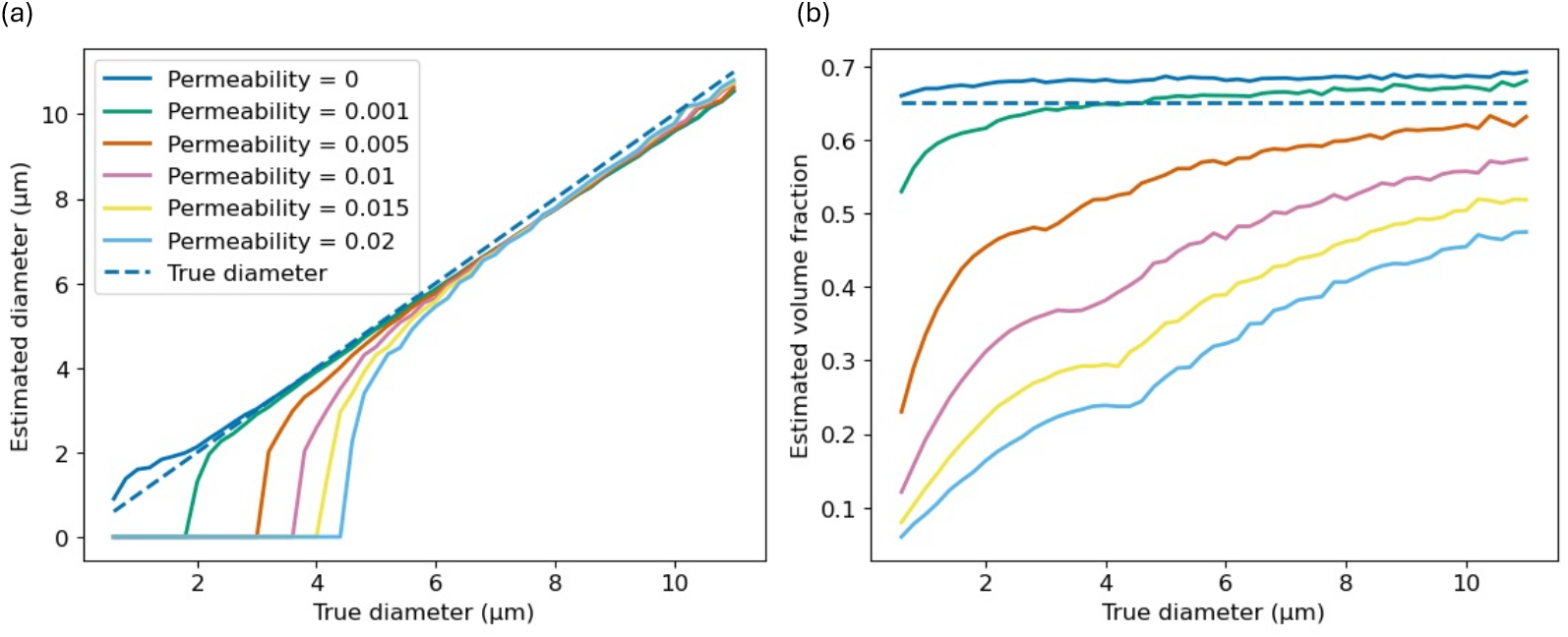
Diameter (a) and volume fraction (b) estimation with semipermeable cylinders, fixed diameter. Permeability caused a reduction in the estimates in both diameter and volume fraction. The diameter estimates additionally exhibit a cutoff behavior, going to zero (the lower bound) when the underlying diameter is below a permeability-dependent cutoff diameter.

When the cylinder diameters became Gamma-distributed, we also observed an underestimation of the cylinder diameter and volume fraction, as shown in Figure 6. The diameter estimation exhibited a similar trend to a single diameter case (orange line in Figure 6a).

**Figure 6.**
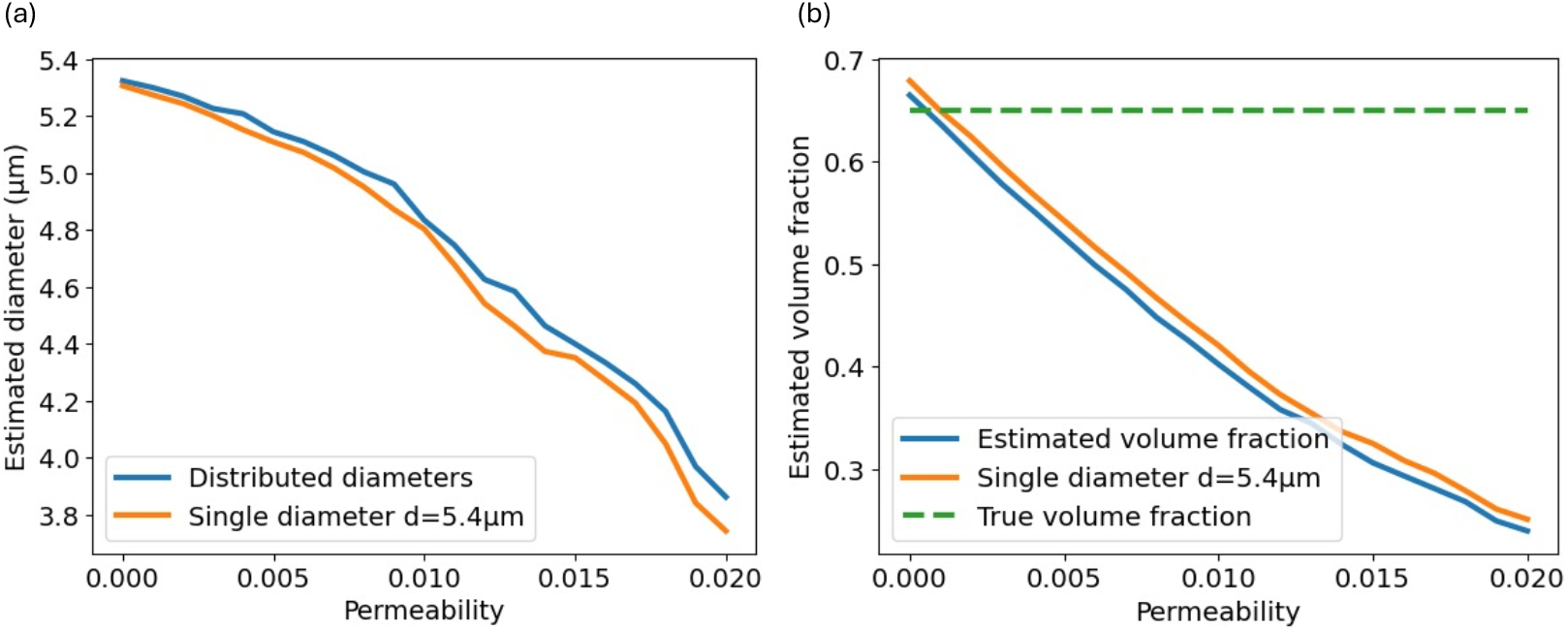
Diameter (a) and volume fraction (b) estimation with semipermeable cylinders, gamma distributed diameter with mean of 4μm and variance of 1μm^2^. The estimate from a single diameter of 5.4μm is plotted to show the similarity in the estimate variation between the distributed diameter case and a uniform diameter case.

### 3.3 Magnetization transfer

Figure 7 plots the estimated diameter and volume fraction as a function of MT strength represented by different effective T_2_ values. The lower the T_2_, the stronger the MT effect is. The figure shows that for the fixed diameter case diameter estimation is broadly independent of MT strength. However, MT led to a large overestimation of the volume fraction across all diameters.

**Figure 7.**
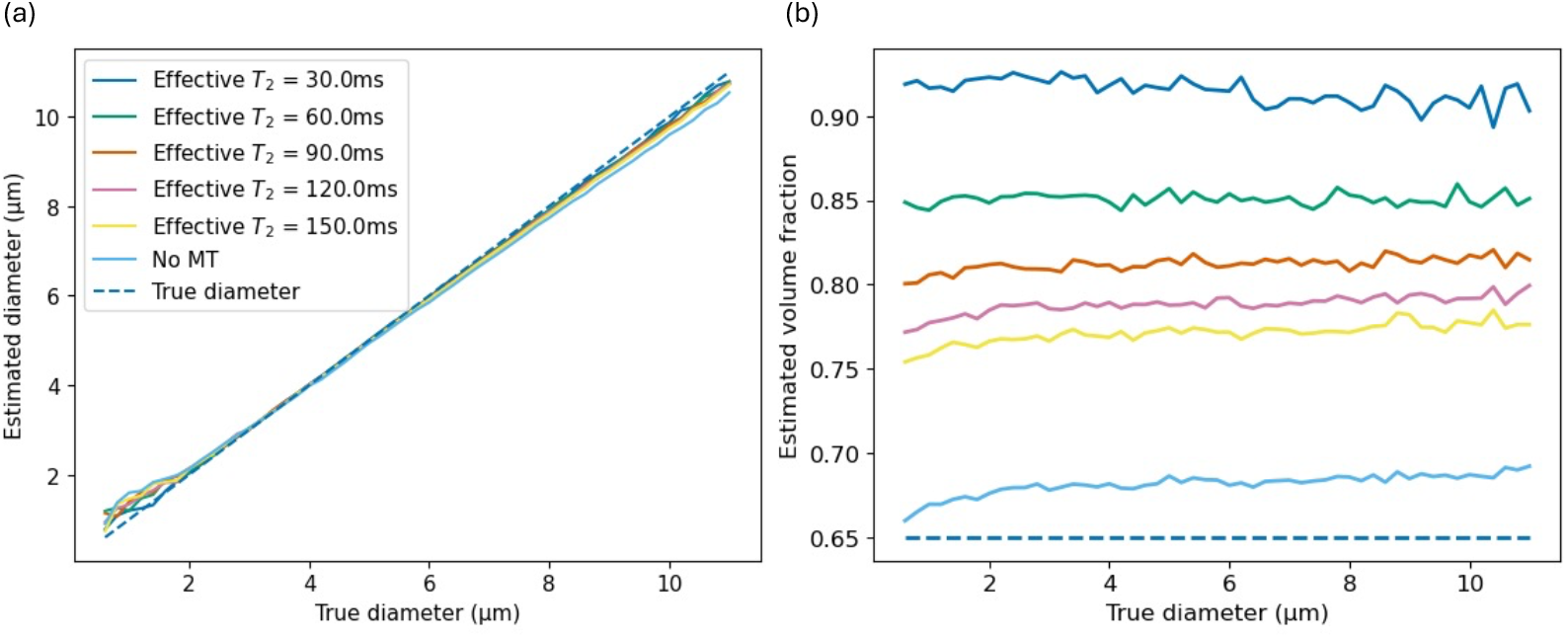
Diameter (7a) and volume fraction (7b) estimation with cylinders with MT, uniform diameter. MT strength had little effect on the diameter estimation but caused large overestimation of the volume fraction. Note that the effective T_2_ is calculated relative to cylinders with a diameter of 2μm.

In the distributed case, we observed an overestimation of both cylinder diameter and volume fraction, as shown in Figure 8. The bias increased as MT strength increased.

**Figure 8.**
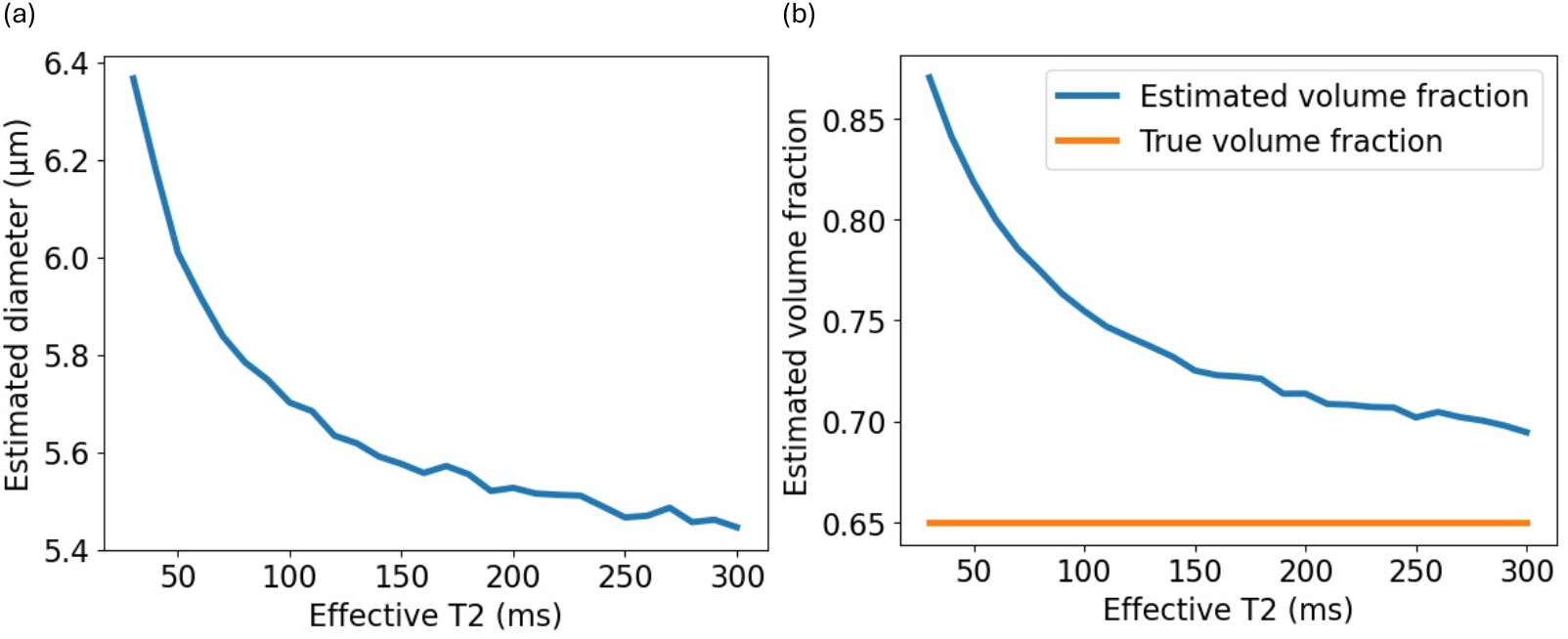
Diameter (8a) and volume fraction (8b) estimation with cylinders with MT, gamma distributed diameter with mean of 4μm and variance of 1μm^2^. MT caused overestimation in both diameter and volume fraction of cylinders.

## 4. Discussion

### 4.1 Permeability effects

The underestimation of the volume fraction observed in Figure 5b has a direct link with permeability. Permeability causes any isochromats that have spent some time in the extra-axonal space to have a displacement profile more similar to an extra-axonal isochromat than an intra-axonal isochromat (see Figure 9). This leads to a large intra-axonal signal reduction under the dMRI encoding and causes the volume fraction to be underestimated in all cases regardless of cylinder diameter. In the fixed diameter case, the underestimation bias is larger for smaller diameters. This is because the macroscopic exchange rate is larger for smaller cylinders for the same permeability value, due to a larger surface-to-volume ratio (see Eqs.2-4 and Figure 2b).

**Figure 9.**
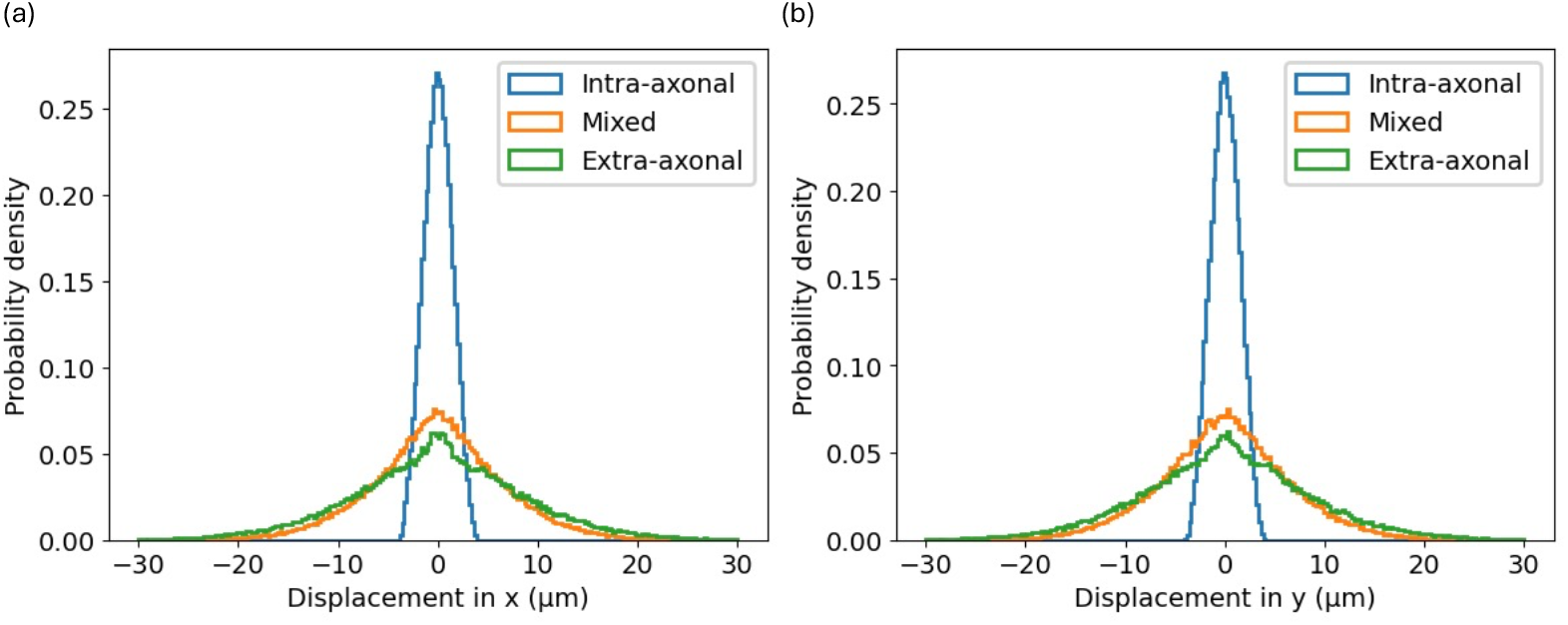
Displacement profile of isochromats along the axes perpendicular to the cylinders. This illustrates isochromats that are within the intra-axonal or extra-axonal space throughout the simulation, and isochromats that spend half of the time in intra-axonal space and half in extra-axonal space (“mixed”). It can be observed that the isochromats that spend half of the time in each compartment have a displacement profile much closer to the extra-axonal displacement profile. This leads to an apparent increase in the extra-axonal signal fraction and underestimation of the axon volume fraction.

The underestimation of the cylinder diameter in semipermeable cylinders (Figure 5a) arises from the construct of the two-compartment model, which associates time-dependent non-Gaussianity in the dMRI signal with the intra-axonal component (Eq.8). When cylinders are impermeable, they restrict the displacement of water inside them, making the dMRI signal deviate further from a Gaussian regime with increasing diffusion time (Δ). This leads to a positive kurtosis (K, an approximate measure of non-Gaussianity) derivative dK/dΔ, with the derivative increasing as the cylinder diameter increases. However, when the cylinder walls are semipermeable, more isochromats cross the cylinder walls with increasing diffusion time and become part of extra-axonal compartment, which has more Gaussian diffusion. This leads to a decrease in dK/dΔ and cancels out the effect of restrictions, which matched similar trends shown in previous studies^56,57^. Consequently, the model fits a smaller cylinder diameter to the data and as the permeability increases the estimate reaches the lower bound – zero, which is observed in Figure 5a. As the increase in dK/dΔ from the restriction is small when the cylinder is small, a weak permeability is enough to cancel it fully and push the diameter estimate down to zero. See Supporting Information 2 for a more detailed kurtosis analysis. The same mechanism also applies to the distributed diameter case which showed similar trend as a fixed diameter case with a diameter larger than the mean of the distribution (see Figure 6a). In addition, this underestimation of restriction size was also observed in a previous study using spherical substrates^58^.

### 4.2 MT effects

When considering MT, the overestimation of volume fraction shown in Figure 7b can be explained by the surface-to-volume ratio differences. Specifically, the intra-axonal compartment has a volume fraction of 0.65, larger than the extra-axonal compartment. Because the intra- and extra-axonal compartment share the same surface area (the cylinder walls), the extra-axonal compartment has a larger surface-to-volume ratio. Since we modelled MT as a surface process, isochromats in the compartment that has a larger surface-to-volume ratio will interact with the cylinder walls more frequently and relax faster. Thus, the extra-axonal signal has a faster effective T_2_ and contributes a smaller fraction of the total signal.

Therefore, the relative fraction of signals that comes from the intra-axonal compartment increases with MT strength and makes the model predict a larger volume fraction estimate. Past literature has also reported that intra-axonal compartment in white matter tends to have a larger effective T_2_ than the extra-axonal compartment^59,60^. We note further that in our simulation the cylinder walls were infinitesimally thin which led to the two compartments having the same surface area. In reality, the axon membrane has finite thickness and myelin sheath thickness will add further to the total thickness. This will make the extra-axonal compartment have a larger surface area than the intra-axonal compartment and amplify the surface-to-volume ratio difference between the two compartments. So, we expect a greater volume fraction overestimation caused by MT in practice.

Further investigations confirmed the volume fraction estimation bias was related to the underlying true volume fraction. When we altered the true volume fraction to 0.5 and 0.35, the overestimation of volume fraction became much smaller (volume fraction of 0.5) or underestimated (volume fraction of 0.35), the opposite effect to the 0.65 volume fraction case (see Figure 10a).

**Figure 10.**
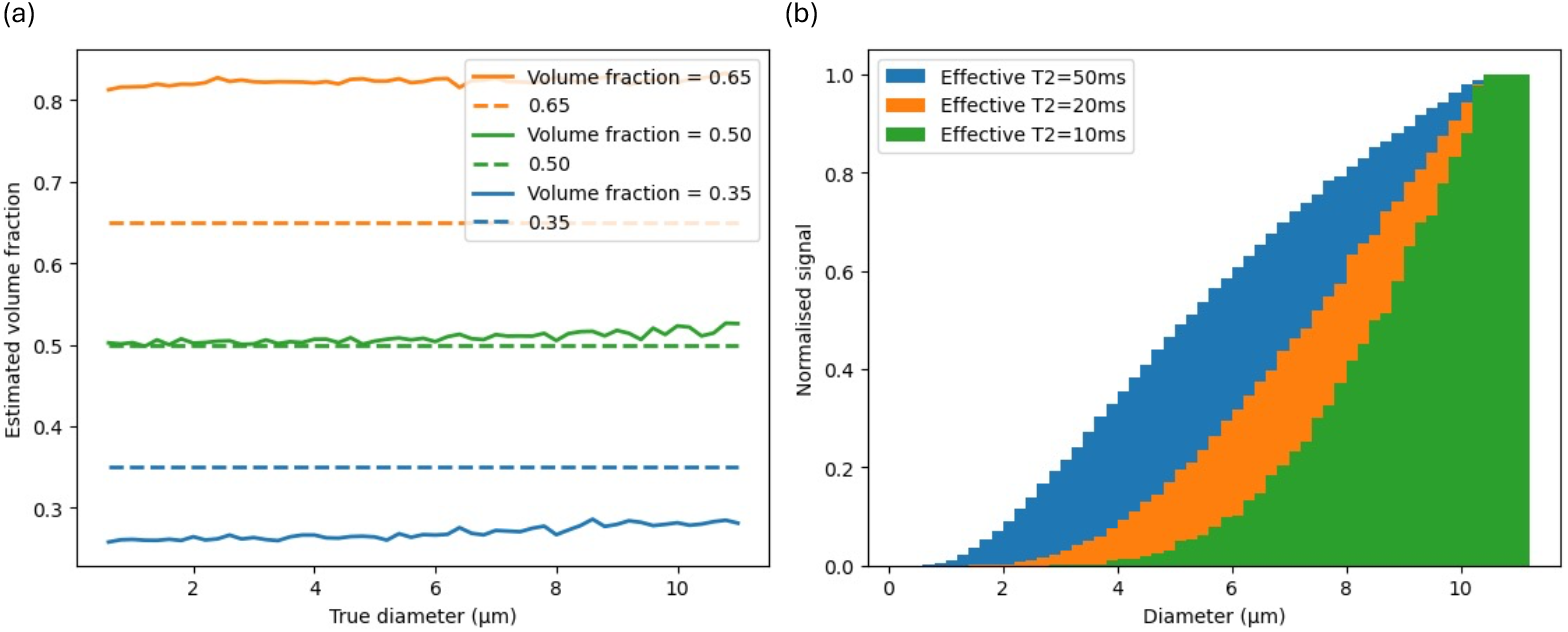
MT results interpretations: Figure 10a shows the volume fraction estimates with different underlying volume fractions (0.35, 0.5, 0.65, marked by the dashed lines). All 3 substrates had the same MT strength (effective T_2_ = 90ms). It can be observed that an underlying volume fraction greater than 0.5 (i.e. 0.65) causes overestimations of volume fraction while an underlying volume fraction smaller than 0.5 (i.e. 0.35) causes underestimations of volume fraction. Figure 10b shows the relative signal levels for axons of different diameters when having the magnetization transfer properties that achieve effective T_2_s of 10, 20 and 50ms for a 4μm-diameter axon.

As for the diameter estimation, since MT doesn’t affect the time-dependence of the kurtosis as permeability does, this explains the minimal change in diameter estimate with MT in fixed diameter cases. MT still affects the kurtosis by altering the relative signal contributions from different compartments but this is captured in the change in the estimated density already.

When the axon diameter became Gamma-distributed, we additionally have to consider the diameter dependence of the macroscopic MT effect. The MT strength is controlled by two diameter-independent parameters – surface density and dwell time (see 2.1.2), with the effective T_2_ dependent on the axon diameter. Since we set the same surface density and dwell time for all cylinders, Eq.5 predicts that smaller axons have a shorter effective T_2_ and larger axons will have a longer T_2_. This matches an experimental observation reported previously^61^. This prediction was verified when we simulated the signal level for a wide range of axon diameters, as shown in Figure 10b. Specifically, we observe a biased weighting in the total signal towards the larger axons as their signal decays more slowly than the smaller axons.

Therefore, during parameter estimation the axon diameter was overestimated as shown in Figure 8a. The overestimation of volume fraction in Figure 8b is still explained by the surface-to-volume ratio difference. However, as more signal came from the axons that had a weaker MT effect, the overestimation bias we observed was also lower compared to the corresponding effective T_2_ in the fixed diameter case (Figure 7b).

### 4.3 Model selection

In this project, we used a simple, two-compartment parallel cylinder model. The motivation was to use a model with well-established properties and limited degeneracies, with variants (e.g. AxCaliber^1^, ActiveAx^62^) widely adopted for microstructural investigations. However, our initial investigation without MT and permeability effects observed a discrepancy between the estimated model parameters (from the two-compartment model) and the true underlying parameters from Monte-Carlo simulations (Figures 4a and b). Further investigation identified that this discrepancy arose from the extracellular space, where the signal model’s assumption of free-Gaussian diffusion was not valid across the diffusion regimes investigated in this study (Figure 4d). However, limited improvement was observed when fitting a time-dependent model^40^ to the extracellular space (Supporting Information 1.2). Typically, the radius bias was a fraction of a micron over the investigated parameter space, with the volume fraction bias <0.05.

### 4.4. Impact

Given these results and interpretations, we argue that MT and permeability can indeed have a non-negligible effect on the dMRI signal. We observed biases in estimated microstructural features (e.g., axon diameter) from the affected dMRI signal using a two-compartment model when MT and permeability are present. More specifically, we noticed that MT and permeability reduce the model’s sensitivity to smaller axons (Figures 5&8). This is because smaller axons have a larger surface-to-volume ratio, which leads to a shorter effective T_2_ with MT and a shorter exchange time with permeability. Furthermore, smaller axons are more likely to be unmyelinated, which leads to even faster exchange through permeability and shorter exchange time. Taken together, MT and permeability’s presence biases dMRI signal further towards the signals generated by the larger axons on top of the previously reported geometry effects^46,63,64^.

Noticeably, in Figure 5, we demonstrate that a small amount of permeability is enough to make a two-compartment model insensitive to axon diameters below 2μm: Specifically, a permeability of 0.001, which translates to an exchange time of ~600ms for axon diameter of 2μm, is enough to cause the estimated diameter to go to zero for axons smaller than 2μm.

Many axons in white matter have diameters below 2μm^1,22,38,40^ and some studies have suggested the white matter exchange time ranges from 150-900ms^65–67^. In addition, for unmyelinated neurons the exchange time measured in grey matter was also shorter than 600ms^46,66,68^. We also demonstrated that MT could bias the axon density estimation through the surface-to-volume ratio difference induced by the underlying volume fractions. The effective T_2_ levels needed to induce a visible bias is comparable to realistic white matter T_2_ values. Our results therefore suggest realistic and biologically relevant MT strength and permeability values can bias dMRI estimates. Thus, dMRI modelling methods that incorporated permeability and multicompartmental relaxation should be considered to produce more accurate estimation of microstructural parameters. No white matter model currently incorporates non-Gaussian diffusion, permeability and MT.

We performed our investigations using Monte-Carlo simulations, observing quantitative effects of MT and permeability on dMRI microstructural estimates. Monte-Carlo simulations allowed us to characterise these effects without having to define complicated analytical equations. Further investigations of more complex geometries are also possible without extra analytical derivations. We have made the code and example data available online at https://github.com/zhiyuzheng1769/MT-and-permeability-effect-on-two-compartment-dMRI-WM-model/tree/main for readers who may be interested in exploring these findings further.

In our investigations we chose a two-compartment model of dMRI signals which models the intra-axonal signal using a parallel cylinder model. As the effect of MT and permeability is only coupled with the surface-to-volume ratio and not the geometry type, our observations should generalise to other dMRI signal models using different geometries (e.g., spheres). In addition, many dMRI white matter signal models share the same two-compartment framework with the one we used and do not account for MT or permeability. Therefore, our findings should also generalise to a wider group of axon diameter mapping methods and substrates.

## Conclusion

We used Monte-Carlo simulations to investigate the effect of MT and membrane permeability on the dMRI signal. Using a substrate consisting of parallel cylinder obstructions, both MT and permeability caused observable changes in dMRI signal and biased microstructural estimates based on a two-compartment model. The changes showed similar trends in both fixed and distributed diameter cases, and are expected to be generalisable to other white matter dMRI signal models and other substrates. Findings demonstrate that Monte-Carlo simulations can provide insights into how MT and permeability can impact dMRI signals, complementing analytical approaches.

## Appendix

### Mathematical relationship between simulator MT variables and effective T_2_

To convert between the simulator MT variables (dwell time and surface density) and the effective T_2_, we need to account for the surface-to-volume ratio (S/V), which is the surface area of the geometry (S) divided by the volume of interest (V). The effective T_2_ can then be derived from the rate of isochromats leaving the free pool (*r*_*out*_) as the isochromats effectively lose their transverse magnetisation instantly when they transfer from the free to the bound pool:

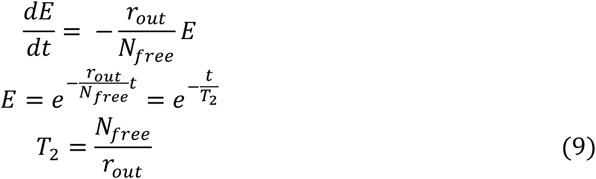

where *E* is the signal attenuation, t is time, and *N*_*free*_ is the number of isochromats in the free pool. The rate of leaving is the same as the rate of entering the free pool (leaving the bound pool) at equilibrium. The rate of entering (*r*_*in*_) is defined as:

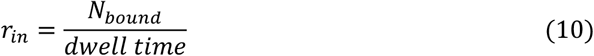

where N_bound_ is the number of isochromats in the bound pool and dwell time is the average time it takes for a bounded isochromat to be released back to free pool. As *r*_*in*_ = *r*_*out*_, we can combine Eq.7 and Eq.8:

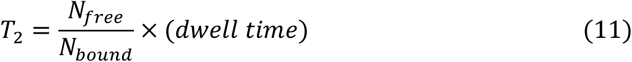

Recall that the surface density is the ratio between surface isochromat density on the obstruction and the volume isochromat density in the free pool. Therefore:

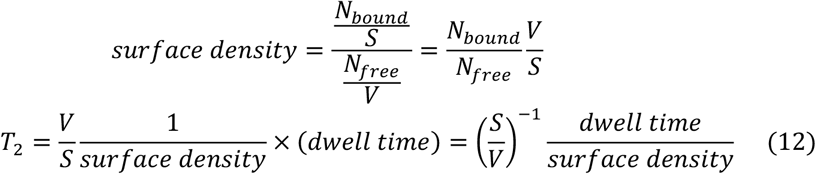

For cylinders:

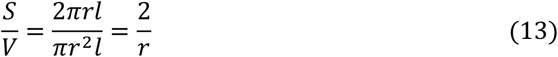

where r is the radius, l is the length. Therefore, we can use Eq.12 (Eq.5 in 2.1.2.2) to get effective T_2_ for the intra-axonal water directly from the dwell time and surface density values.

## Acknowledgements

ZZ is supported by University of Oxford and China Scholarship Council. KLM is supported by a Wellcome Trust Senior Research Fellowship (224573/Z/21/Z). BCT is supported by a Wellcome Trust Sir Henry Wellcome Fellowship (222829/Z/21/Z). MC is supported by a Wellcome Trust Collaborative Award (215573/Z/19/Z). The Centre for Integrative Neuroimaging was supported by core funding from the Wellcome Trust (203139/Z/16/Z and 203139/A/16/Z). This research was funded in whole, or in part, by the Wellcome Trust (224573/Z/21/Z, 222829/Z/21/Z, 215573/Z/19/Z, 203139/Z/16/Z, 203139/A/16/Z). For the purpose of open access, the author has applied a CC BY public copyright license to any Author Accepted Manuscript version arising from this submission.

