## Supporting Information for "Investigating the sensitivity of the diffusion MRI signal to magnetization transfer and permeability via Monte-Carlo simulations"

### 1 Selection of compartment models in the two-compartment model

#### 1.1 Evaluating analytical and simulation-based models for the intra-axonal signal

To accurately model the intra-axonal signal, we used MCMRSimulator to generate a dictionary of intra-axonal signals from parallel cylinders with different diameters at a spacing of 0.2µm. We then constructed a projection from cylinder diameter to the intra-axonal signal by interpolating between the discrete value pairs in the simulated dictionary. This allowed us to leverage the Monte-Carlo simulation which provides the ground truth for our modelling and make fewer assumptions versus existing analytical models (e.g. Callaghan[1], Van Gelderen[2]) which are based on the Gaussian Phase Approximation. As shown in Figure S1, breakdown of these assumptions in restricted substrates leads to deviation in the estimated cylinder diameter, while our interpolated dictionary approach gave accurate estimates.

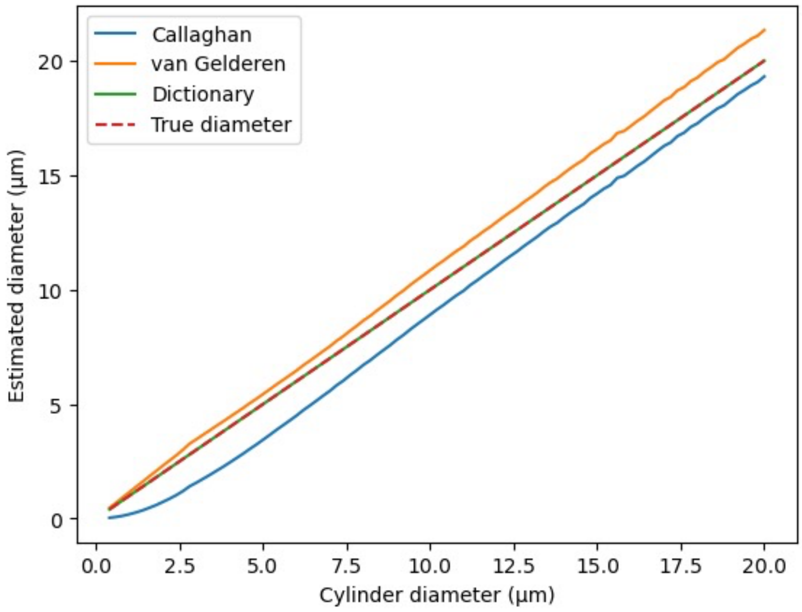


Figure S1: **Intra-axonal signal model selection**: Figure S1 shows the diameter estimate using the intra-axonal signal with different signal models. Both Callaghan and van Gelderen models' (which are based on the Gaussian phase approximation) estimations deviated from the ground truth generated by Monte-Carlo simulation.

#### 1.2 Evaluating time dependence and kurtosis models to characterize the extra-axonal signal

To effectively characterize the extra-axonal signal, we performed comparisons using a (1) Gaussian, (2) time-dependent and (3) kurtosis model. Specifically, the Gaussian model was implemented as:

$$E_e = e^{-bD_e}$$

where $E_e$ is the extra-axonal signal attenuation, $b$ is the b-value of the applied sequence, $D_e$ is the diffusivity of the extra-axonal space.

The time dependent model was implemented as:

$$E_e = e^{-b\left(D_\infty + A\left(\frac{\ln\left(\frac{\Delta}{\delta}\right)+\frac{3}{2}}{\Delta - \frac{\delta}{3}}\right)\right)}$$

where $D_\infty$ is the bulk diffusivity, $A$ is a characteristic coefficient, $\Delta$ is the diffusion time, $\delta$ is the gradient duration. This implementation was initially proposed by De Santis et al.[3].

And the kurtosis model was implemented as:

$$E_e = e^{-bD_e + \frac{1}{6}b^2 D_e^2 K}$$

where K is the kurtosis.

As shown in Figure S2, we found that the three models produced similar trends across the investigated diameter and (intra-axonal) volume fraction regimes. Specifically, the Gaussian and kurtosis models yielded almost identical results. The time-dependent model yielded some improvement in diameter estimation in small diameter regimes, and volume fraction in high diameter regimes but its estimates remained noisy.

Given the small differences between the three extra-axonal models and the additional parameters required for the time-dependent or kurtosis model, we proceeded with the Gaussian diffusion model. This approach is consistent with many existing two-compartment models of white matter and facilitates characterisation of the extra-cellular signal compartment.

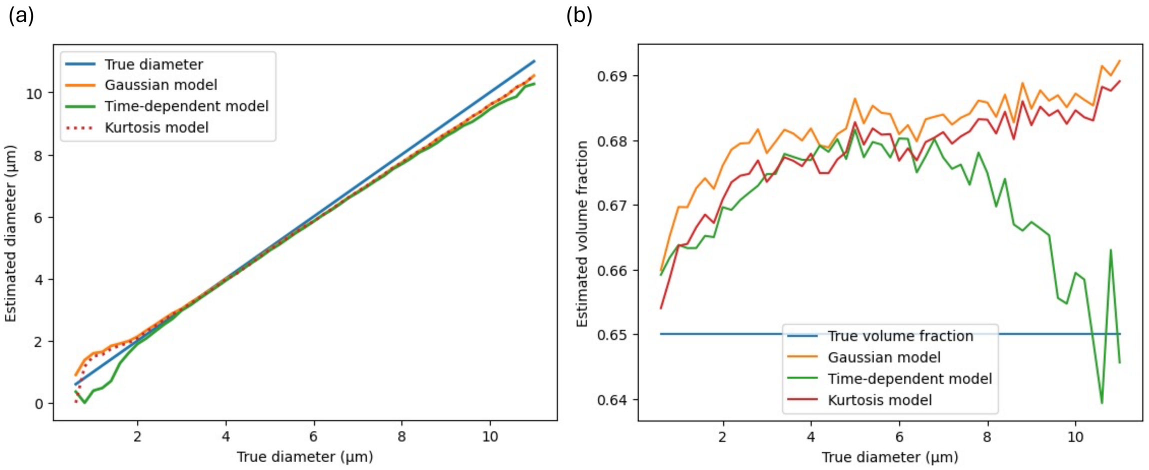


Figure S2: **Extra-axonal signal model selection**: Figures S2a&S2b show Axon diameter and volume fraction estimates without permeability and MT using different extra-axonal signal models in the fixed diameter case. The performance of the kurtosis model is very similar to the Gaussian model. The time-dependent model outperformed the Gaussian model in the volume fraction estimation of large cylinders and diameter estimation of small cylinders.

### 2 Kurtosis quantification for semi-permeable cylinders

Our two-compartment model associates any time-dependent non-Gaussianity in the dMRI signal with the intra-axonal component and use it to estimate the cylinder diameter. To understand why we observed an underestimation of cylinder diameter in the semi-permeable cylinders, we quantified the non-Gaussianity in the simulated diffusion-weighted signal by estimating its kurtosis. Kurtosis K is defined in a second order correction for the Gaussian diffusion signal model to capture the non-Gaussianity:

$$\frac{S}{S_0} = e^{-bD+\frac{1}{6}b^2D^2K}$$

Where $S$ is the diffusion-weighted signal, $S_0$ is the non-diffusion-weighted signal, $b$ is the b value, $D$ is the apparent diffusivity. To estimate the kurtosis, we fitted a quadratic function of the b-value to the natural logarithm of the diffusion-weighted signal attenuation $\frac{S}{S_0}$. The coefficient of the second order term is then $\frac{1}{6}D^2K$ and D can be directly obtained as the coefficient of the first order term.

Figure S3a displays the kurtosis estimates for impermeable cylinders at different diffusion times (Δ) as a function of cylinder diameter. As the diameter increases, we observe an increase in the time-derivative of the kurtosis dK/dΔ arising from the restriction size effect: at a diameter around 2μm, kurtosis only increases less than 0.1 as the diffusion time increases from 10 to 40ms, whereas the same increase in diffusion time caused an increase of 0.4 in kurtosis when the diameter is 8μm.

Figure S3b displays the kurtosis estimates at different diffusion times as a function of permeability for a fixed cylinder diameter (4μm). As the permeability increases, dK/dΔ starts decreasing, displaying the opposite trend to Figure S3a. Specifically, the kurtosis at shorter diffusion times is smaller than the kurtosis at longer diffusion times for low permeability, with the opposite trend at high permeability. Taken together, when incorporating permeability, the relationship between diffusion time and kurtosis amplitude becomes more reflective of smaller diameters in the impermeable cylinder case (Figure S3a), leading to the underestimation of diameter when using a two-compartment model.

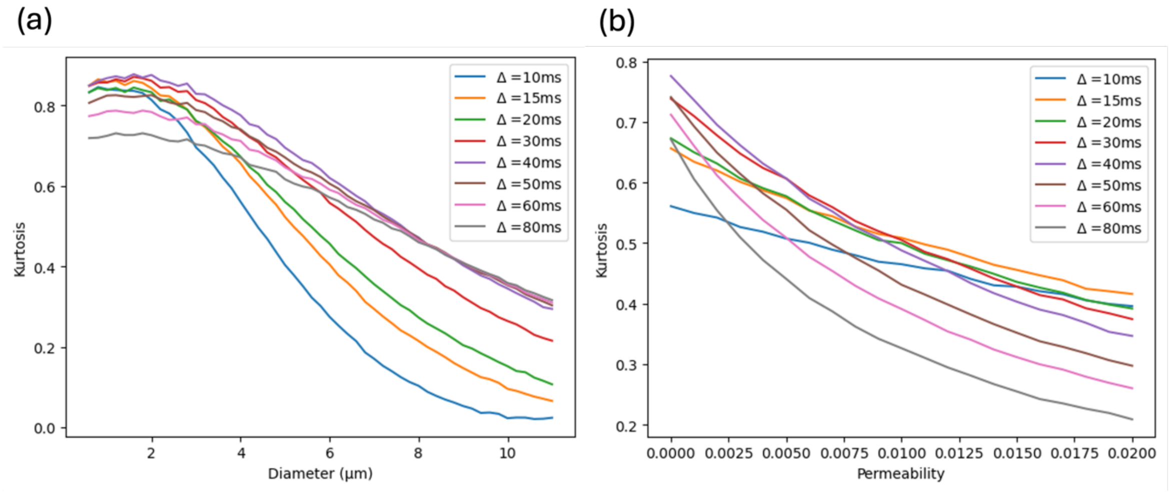


Figure S3: **Kurtosis variation with diffusion time, cylinder diameter and permeability** Figure S3a shows the kurtosis at different diffusion times across different diameter with impermeable cylinders and Figure S3b shows the kurtosis at different diffusion times across different permeability for cylinders with a fixed diameter of 4μm.
